# Proton Translocation and Q-Cycle Efficiency at the Cytochrome *b_6_f* Stromal Q_n_ Site Require Conserved Subunit-IV Residue Asp35 in *Chlamydomonas reinhardtii*

**DOI:** 10.64898/2026.11.20.768064

**Authors:** Simon Bütfering, Yuval Milrad, Rafał Pietras, Shi-Ichiro Ozawa, Felix Buchert, Marcin Sarewicz, Martin Scholz, Artur Osyczka, Michael Hippler

**Affiliations:** Institute of Plant Biology and Biotechnology, University of Münster, Münster 48143, Germany; Faculty of Biochemistry, Biophysics and Biotechnology, Department of Molecular Biophysics, Jagiellonian University, Kraków, Poland; Institute of Plant Science and Resources, Okayama University, Kurashiki, Okayama 710-0046, Japan

## Abstract

Oxygenic photosynthesis relies on the efficient operation of the cytochrome *b*_*6*_*f* complex (*b*_*6*_*f)*, which acts as the central pacemaker linking linear and cyclic electron transfer chains *via* its catalytic Q-cycle. Unlike its mitochondrial counterpart, the chloroplast *b*_*6*_*f* features a unique heme *c*_*n*_/*b*_*n*_ pair and an intricate network of conserved water molecules and polar residues near the stromal Q_n_ site, implicated in proton translocation and plastoquinone (PQ) reduction. Here, we investigate the functional significance of the conserved aspartate residue of subunit-IV (PetD:D35), which is located in the centre of the stromal proton resupply channel. Using chloroplast site-directed mutagenesis in *Chlamydomonas reinhardtii*, we generated and characterized three substitution mutants to: glutamate, serine and alanine, to test the contributions of side-chain length, charge, and polarity. Biochemical isolation and EPR spectroscopy revealed that the assembly of the *b*_*6*_*f* and the coordination of the Rieske-ISP subunit remain uncompromised across all isolated *b*_*6*_*f* variants. However, flash-induced single-turnover kinetics and electrochromic shift measurements demonstrated that the substitution to alanine creates a severe kinetic blockade at the Q_n_ site, trapping low-potential *b*-hemes in a reduced state and crippling forward electron transfer through the high-potential chain. These findings indicate that PetD:D35 is essential for proton resupply and Q-cycle efficiency, linking stromal Q_n_ site dynamics to broader redox regulation and photosynthetic viability.

## Introduction

Oxygenic photosynthesis fundamentally shapes the biosphere by converting solar radiation into chemical energy, driving primary production and sustaining atmospheric composition. This process relies on a continuous electron transfer chain embedded within the thylakoid membrane, where light energy absorbed by photosystem II (PSII) and photosystem I (PSI) is coupled to transmembrane proton translocation (Nelson, 2011; Rutherford et al., 2012). Connecting the two photosystems, the cytochrome *b*_*6*_*f* complex (*b*_*6*_*f*) acts as the central rate-limiting component and orchestrator of photosynthetic electron flow (Malone et al., 2021; Sarewicz et al., 2021). Through its catalytic Q-cycle mechanism, the complex generates a trans-membranal proton translocation, enhancing the proton motive force required for ATP synthesis. This process happens *via* bifurcation of two electron transfer chains, where the plastoquinol (PQH_2_) is being oxidized at the luminal Q_p_ site (Cramer et al., 2011). From there, electrons are channelled towards Cyt*f via* the high-potential chain, and towards a plastoquinone (PQ) at the stromal Q_n_ site *via* the low-potential chain (Figure.1A). This site is located on the stromal n-side of the thylakoid-membrane embedded *b*_*6*_*f*. A comparison between *b*_*6*_*f* complex with its mitochondrial homologue, cytochrome *bc*_*1*_ (*bc*_*1*_) complex (Hasan et al., 2013; Szwalec et al., 2022), reveals a unique heme situated within the Q_n_ site, named *c*_*n*_. This heme lays in close proximity to heme *b*_*n*_ (Kurisu et al., 2003; Stroebel et al., 2003), that is found in *bc*_*1*_. This additional heme *c*_*n*_ occupies the ubiquinone binding niche of the *bc*_*1*_ complexes (Kurisu et al., 2003; Pintscher et al., 2024), and is covalently bound to just one cysteine 35 of cytochrome *b*_*6*_. The heme iron does not feature axial ligands typical for hemes in cytochromes: one axial ligand position is occupied by water molecule while the other position remains unoccupied (Twigg et al., 2009). However, the structures of *b*_*6*_*f* with inhibitors revealed that Q_*n*_ site inhibitors can bind at this position fulfilling the role of axial ligand (W_1_, Figure.1B) (Jones and Whitmarsh, 1988; Yamashita et al., 2007; Hasan et al., 2013). The structure of *b*_*6*_*f* frozen during the catalytic turnover revealed that PQ analogue also binds in the similar niche, but unlike inhibitors, does not ligate the heme (Pintscher et al., 2024). The position of the PQ within this region is seemingly confined by heme *c*_*n*_ and arginine 207 of subunit *b*_*6*_ (Cyt*b*_*6*_:R207, Figure.1B).

**Figure 1.**
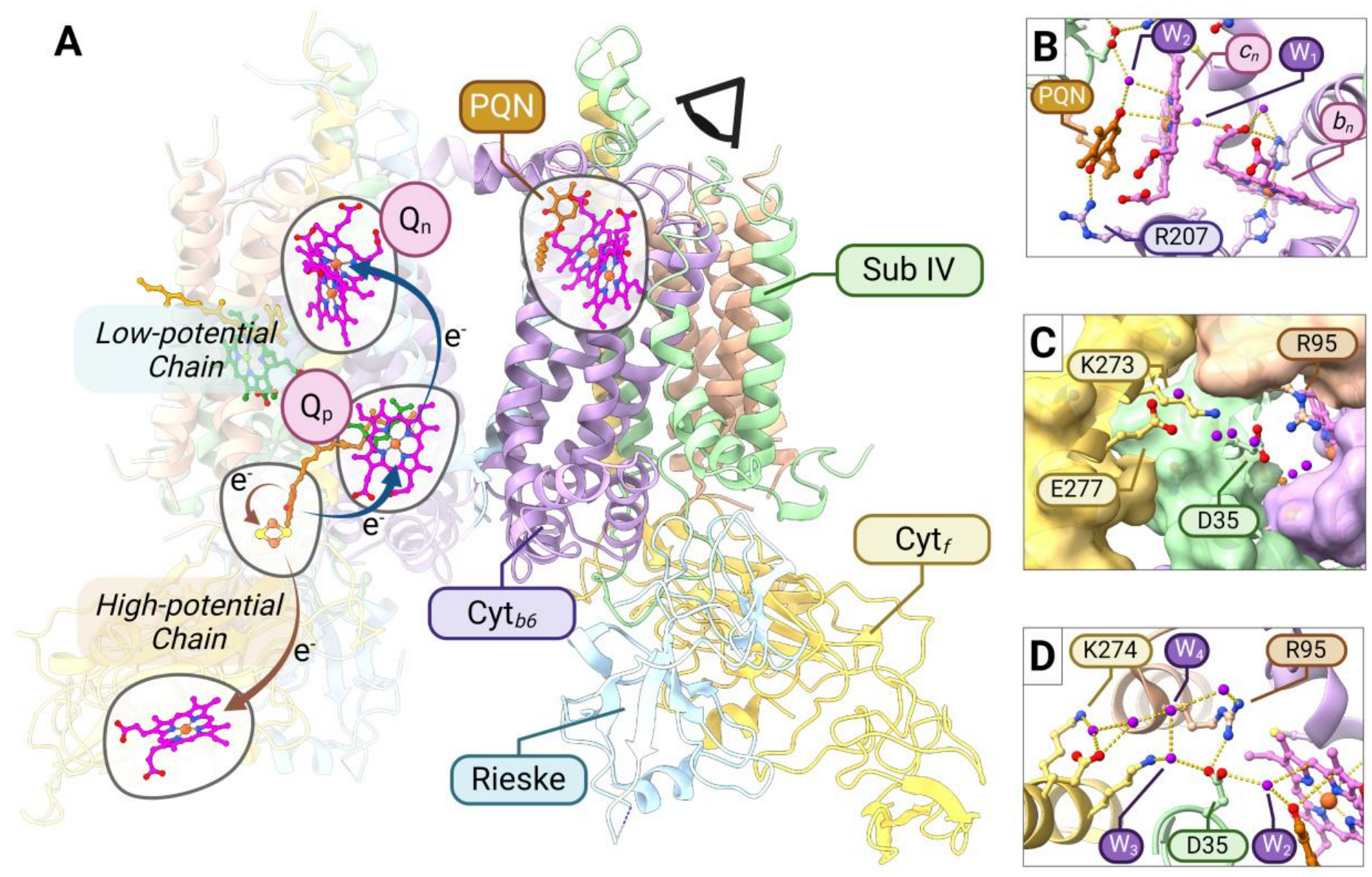
The positioning of the water channels on the stromal side of Subunit IV: Structural model of the dimeric cytochrome *b*_*6*_*f* complex highlighting key cofactors, transmembrane subunits, and internal electron/proton transfer pathways. (**a**) Overview of the complex showing Subunit-IV (green), cytochrome *b*_6_ (lilac), cytochrome *f* (yellow), Rieske-ISP (cyan) and the supporting helices (Pet;G,L,M,N in peach). The panel also show the positioning of the hemes (*c*_*n*_, *b*_*n*_, *b*_*p*_, *f* in magenta), the [2Fe-2S] cluster, and plastoquinones (PQN, orange). In addition, the scheme describes the electron channeling on the low-potential chain (towards the hemes *c*_*n*_ and *b*_*n*_ in Q_*n*_) and the high-potential chain (towards heme *f*). (**b–d**) Detailed views of the stromal Q_n_ environment, focusing on; the position of hemes *c*_*n*_ and *b*_*n*_, bridging water (W_1_ and W_2_ in purple) molecules and PQN (**b**), top view of the water channel (**c**), and detailed view on the water channel, which is confined by residues; PetD:D35, PetN:R95, PetA:K273, PetA:K274 and PetA:E277 (**d**). The structure visualization is based on a cytochrome *b*_*6*_*f* structure of Chlamydomonas reinhardtii (PDB:1Q90) where waters and PQN are superposed in accordance with their localization based on Spinacia oleracea (PDB: 9ES8).

Recently, high-resolution structural and biophysical studies have revealed a complex network of water molecules and polar residues near the stromal Q_n_ site, providing critical clues regarding the mechanism of transmembrane proton translocation and quinone reduction (Pintscher et al., 2024, see also Figure.1 C,D). Moreover, studies showed that heme *c*_*n*_ exhibits a pH-dependent redox midpoint potential of approximately ∼60 mV per pH unit, which grants it the ability to undergo deprotonation upon oxidation (Alric et al., 2005). This redox-active heme environment acts in concert with a dedicated proton transfer network, wherein a structurally conserved water molecule (W_2_) forms a bridge between the PQ and a highly conserved aspartate residue on subunit IV(SubIV:D35, Figure.1, see also Supplemental Fig.S1). This aspartate stands in the centre of an intricate water and proton channel of water molecules, formed by; PetD:D35, PetN:R95, PetA:K273, PetA:K274 and PetA:E277 (Figure.1). Such architecture is capable of efficiently protonating reduced PQ, as part of a potential proton resupply pathway. To test the functional significance of this putative proton and water channel network, this work examines the role of the conserved PetD:D35 residue in PQ reduction and Q_n_ site turnover. Using site-directed mutagenesis in the green alga *Chlamydomonas reinhardtii*, we generated and characterized three distinct chloroplast substitution mutants; glutamic acid (D35E), serine (D35S), and alanine (D35A), to test the contributions of side-chain length, polarity, and charge. Combining biochemical isolation with pulsed-amplitude modulation (PAM) fluorometry, low temperature chlorophyll *a* fluorescence, and Joliot-type spectroscopy (JTS), our findings implicate important contribution of PetD:D35 to efficient proton re-supply, substrate coordination, and Q-cycle efficiency, offering broader implications for how stromal Q_n_ dynamics influence linear and cyclic electron flow and regulate STT7 activation.

## Results

### Mutant Generation and Cytochrome *b*_6_*f* Complex Integrity

To investigate the functional role of PetD:D35, which is located in the stromal water channel (Figure.1), we generated three lines of mutants (D35A, D35S and D35E). These site-directed strains were successfully generated *via* biolistic transformation to the chloroplast. To enable complex targeted purification, the vectors were introduced to a petA-HisX6 parental strain (see Milrad et al., 2025). Homoplasmy and successful insertion were confirmed *via* sequencing analysis of two independent clones per substitution (Supplemental Fig.S2). A resistance-cassette containing wild-type strain (WT_cas_) was used as a control across all experiments. To test the expression level of *b*_*6*_*f* genes, cells were cultured, digested and subjected to a mass spectrometry-based label-free protein quantification (LFQ). Results were normalized based on the averaged values gained for ATPase subunit (an average of; atpA, atpB, ATPC, ATPD, atpE and atpF), and assembled in box plots, comparing each subunit of *b*_*6*_*f* (PetA, PetB, PETC, PetD and PETM, see supplementary Fig.S3). According to our data, there seem to be no significant difference between any of the strains, regarding the expression of *b*_*6*_*f* subunits. To confirm the structural assembly and stability of the mutant *b*_*6*_*f* complexes, complexes were purified and loaded on an SDS-PAGE. Immunoblotting and heme-staining assays were performed. Sucrose density gradient isolations yielded intact dimeric *b*_*6*_*f* complexes for all strains (Figure.2A). Ponceau S loading controls and AtpB immunoblots of isolated thylakoid membranes indicated equal total protein loading, whereas PetD immunoblotting revealed slightly lower signal intensities for the D35S and D35A variants compared to WT and D35E (Figure.2A). This suggests potential minor reductions in PetD accumulation or increased susceptibility to protease-mediated remodelling. In-gel TMBZ peroxidase staining of non-reducing SDS-PAGE gels confirmed the presence of high-spin hemes *f* and *c*_*n*_ across all strains, with staining intensities scaling with the corresponding Coomassie-stained protein levels of subunits PetA and PetB (Figure.2A). Furthermore, low-temperature X-band CW-EPR spectroscopy of isolated complexes verified that the intrinsic [2Fe-2S] cluster of the Rieske iron-sulfur protein remain structurally uncompromised across all variants (Figure.2B).

**Figure 2.**
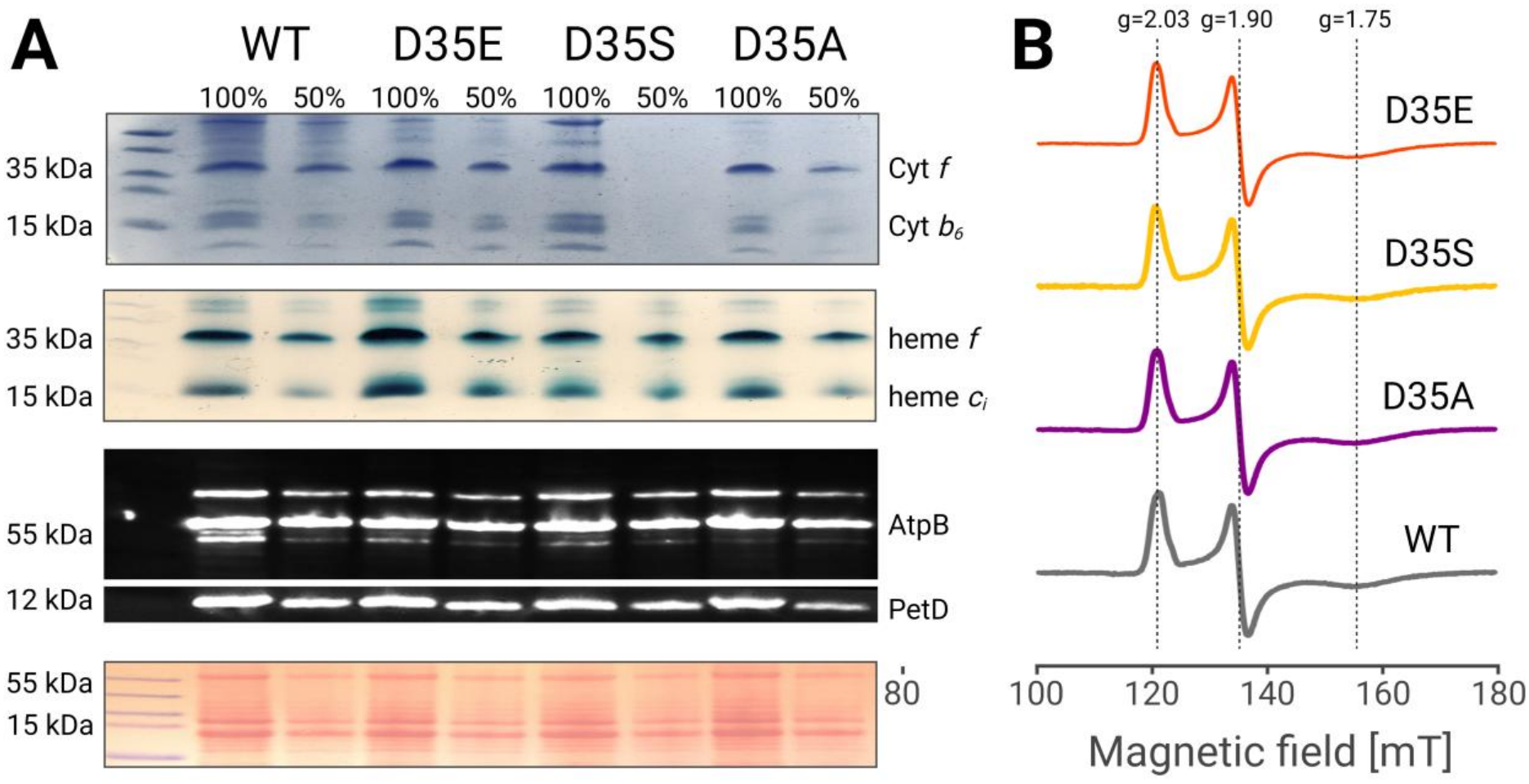
Biochemical evaluation of isolated cytochrome *b*_6_*f* complexes: *b*_*6*_*f* from the WT and mutant variants (D35E, D35S and D35A) were purified and their stability was tested. (**A**) *in-gel* visualizing of the high-spin hemes *f* and *c*_*n*_. The panel presents (from top) Coomassie blue staining of the SDS-PAGE, TMBZ peroxidase staining, immunoblots of isolated thylakoid membranes probed with AtpB and PetD antibodies and Ponceau stains for loading control. (**b**) Baseline-corrected X-band CW-EPR spectra of the Rieske [2Fe-2S] cluster region demonstrating structural retention across all strainsg NiNTA column. (**A**) Coomassie blue-stained SDS-PAGE and in-gel TMBZ peroxidase staining visualizing high-spin hemes *f* and *c*_*n*_ for WT and mutant variants (D35A, D35S and D35E). Immunoblots of isolated thylakoid membranes probed with AtpB and PetD antibodies alongside Ponceau stains for loading control. (**B**) Baseline-corrected X-band CW-EPR spectra of the Rieske [2Fe-2S] cluster across all strains.

### Physiological Characterization and Photoautotrophic Growth

The physiological consequences of the PetD:D35 substitutions were evaluated via photoautotrophic growth assays on minimal medium under low (50 µE m^-2^ s^-1^) and high-light (225 µE m^-2^ s^-1^) conditions. While the D35E substitution exhibited a WT like growth phenotype, the D35S mutant showed a moderate growth deficit, and the D35A mutant displayed severe photoautotrophic growth impairment under both light regimes (Figure. 3A). Pulsed-amplitude modulation (PAM) fluorometry revealed a rapid, early-stage decline in the effective quantum yield of Photosystem II (Y_II_) for D35S and D35A under increasing actinic light intensities (PAR), with D35A dropping sharply at low thresholds (35-80 µE m^-2^ s^-1^, Figure.3B). Correspondingly, the reduction state of the PSII acceptor side (1-qP) increases in D35S and D35A, indicating pronounced over-reduction of the intermembrane PQ pool even under moderate illumination. Despite these redox imbalances, non-photochemical quenching (NPQ) induction kinetics remained comparable among the strains, and 77 K chlorophyll *a* fluorescence spectrum confirmed that all mutants retain full capacity for state transitions (Supplemental Fig.S4). Joliot-type spectroscopy (JTS) measurements of steady-state electron transfer rates (ETR) under aerobic and anoxic conditions demonstrated a significant decrease in steady-state ETR for the D35A mutant, whereas D35E maintained wild-type rates (Figure.3 C, D). Measurements done under anoxia, which favours cyclic electron flow (CET), showed similar relative trends, underlining a general deceleration of photosynthetic electron transport originating from the mutated *b*_*6*_*f* complex.

**Figure 3.**
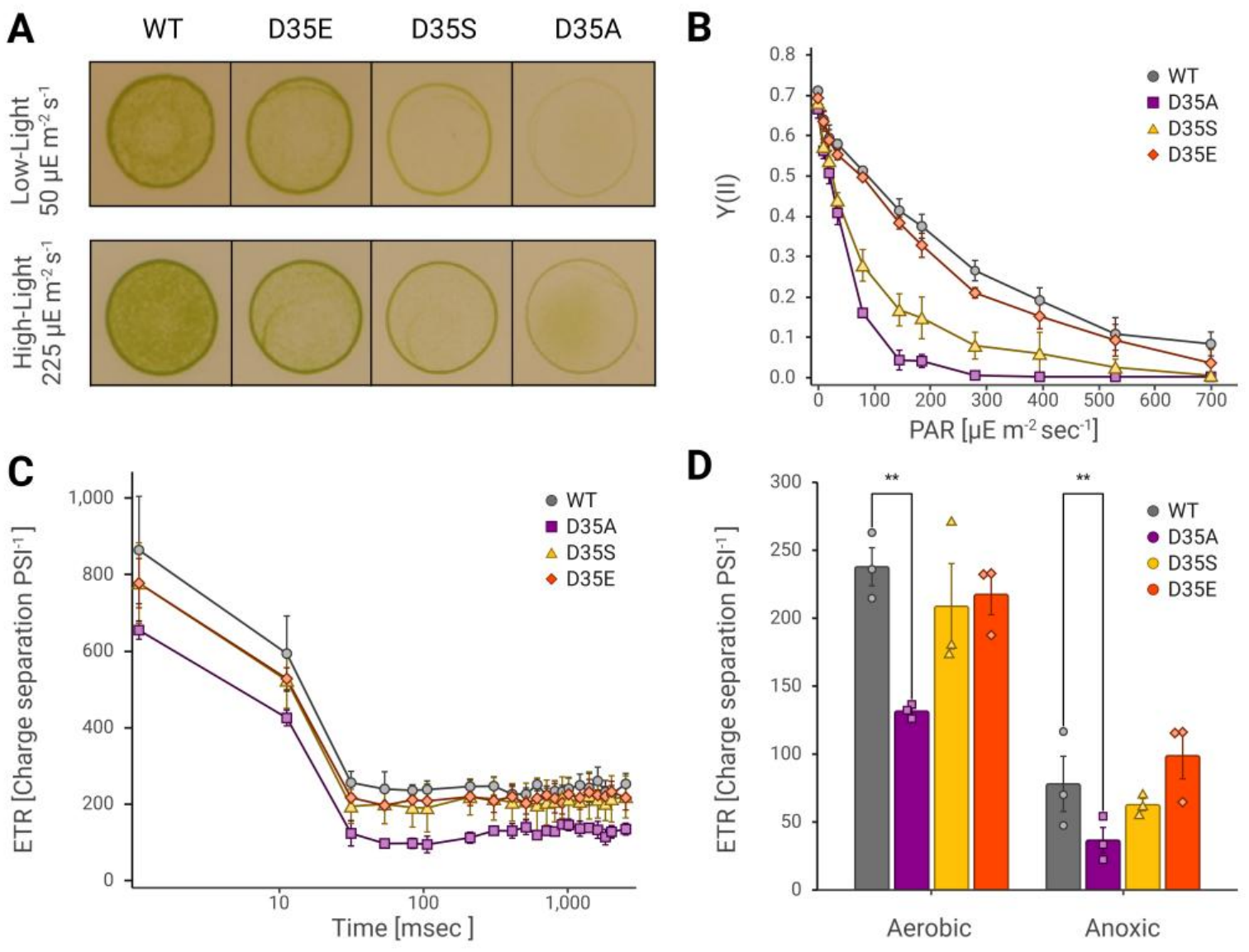
Physiological and photosynthetic characterization of PetD:D35 mutants: (**a**) Spot-test growth assays on minimal (TP) medium under low (50 µE m^-2^ s^-1^) and high (225 µE m^-2^ s^-1^) light. (**b**) PSII quantum yields over irradiance curves were measured *via* PAM chlorophyll *a* fluorometry. (**c**) Electron transfer rates (ETR) determined *via* DIRK measurements. This scheme was conducted under both aerobic and anoxic conditions. (**d**) Steady-state ETR values were averaged for all conditions. In-condition group comparison was made using a two-way ANOVA followed by Holm–Šídák’s multiple comparisons test to compare each mutant with WTAsterisks (**) denote statistical significance at p < 0.01. Each group consisted of three biological replicates (n = 3). Figure assembly and statistic tests were generated using Biorender.com.

### P700 Redox Kinetics and PSI Limitations

The redox kinetics of Photosystem I special pairs (P700) were monitored via absorbance changes at 705 nm (with a reference line at 740 nm) during actinic illumination and subsequent darkness (Figure.4A). In WT and D35E strains, illumination triggered a biphasic P700 oxidation pattern. In contrast, D35A exhibited an accelerated initial oxidation transient, coupled with a significantly elevated fraction of constantly oxidized P700 (Y_ND_) under steady-state conditions (Figure.4B). This pronounced donor-side limitation likely reflects a bottleneck in the resupply of reduced plastocyanin by the upstream *b*_*6*_*f* complex. Under anoxic conditions with PSII inhibitors (DCMU+HA), D35A displayed an extreme oxidation overshoot, pointing to severe acceptor- and donor-side redox constraints. Moreover, we tracked electrochromic shifts (ECS) of the flash-induced cultures from our anoxic treated samples by measuring absorbance changes at 520 nm (with a reference line at 546 nm, Bailleul et al., 2010; Buchert et al., 2020). The results were plotted on a logarithmic scale, where the trace was normalized using the a-phase as one PSI charge separation. The increase of the b-phase was observed in the WT and S35E cultures for up to 20 msec post laser flash (Figure.5A). Interestingly, D35S showed a diminished increase, while D35 was absent from a b-phase increase completely. Following the initial increase, the traces transitioned into the c-phase, where the ATP-synthase consumes the generated PMF and overwrites the PQ pool dependent *b*_*6*_*f* contribution to ECS in the dark. This indicates a diminished electron transfer efficiency of the *b*_*6*_*f* complex, in accordance with our previous described observations.

**Figure 4.**
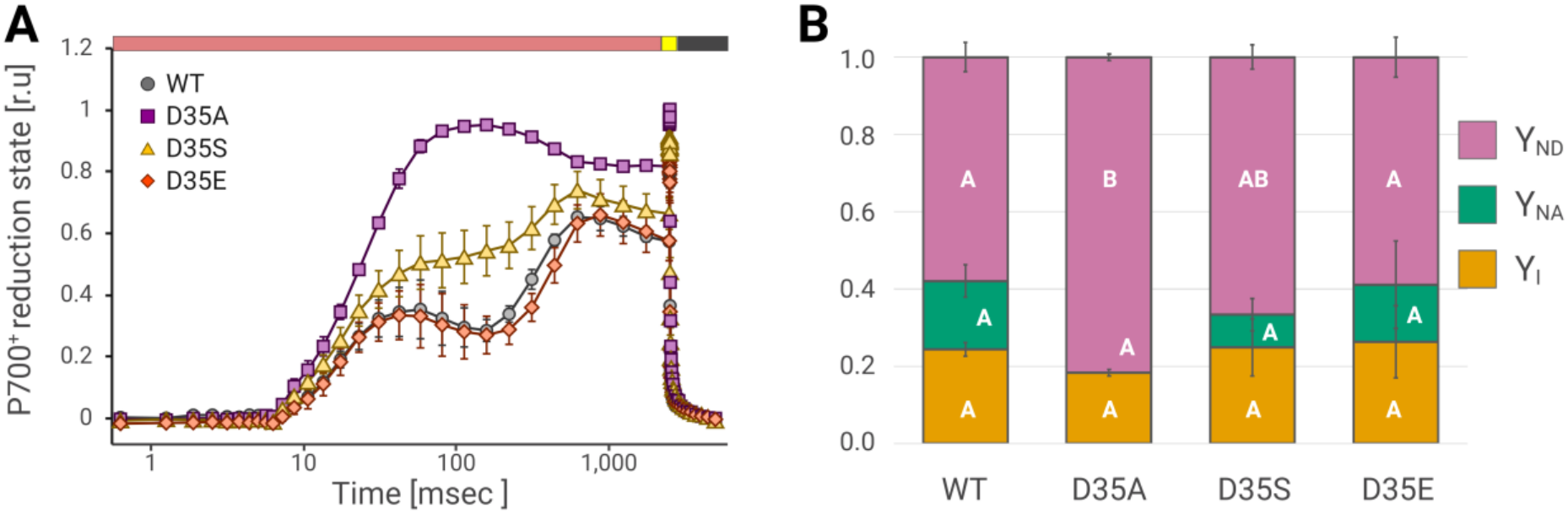
Photosystem I donor-side limitations in accordance to the functionality of the water channels on SubIV: (**a**) Normalized P700^+^ redox kinetics during 2.5 s actinic illumination (600 µE m^-2^ s^-1^, X-axis red bar), followed by a 20 ms of saturating pulse (3000 µE m^-2^ s^-1^ X-axis, yellow bar), and dark relaxation (black bar). (**b**) Partitioning of PSI quantum yield (Y_I_-pink), acceptor-side limitation (Y_NA_-green), and donor-side limitation (Y_ND_-gold) derived from saturation pulse analysis. *in-condition* group comparison was made using a two-way ANOVA-Holm-Šidák test with respect to WT. Significant difference between mutants (WT, D35E, D35S, D35A) is grouped and indicated by letters. Each group is consistent of three biological replicates (n=3) with p.v < 0.01. Figure assembly and statistic tests were generated using Biorender.com.

**Figure 5.**
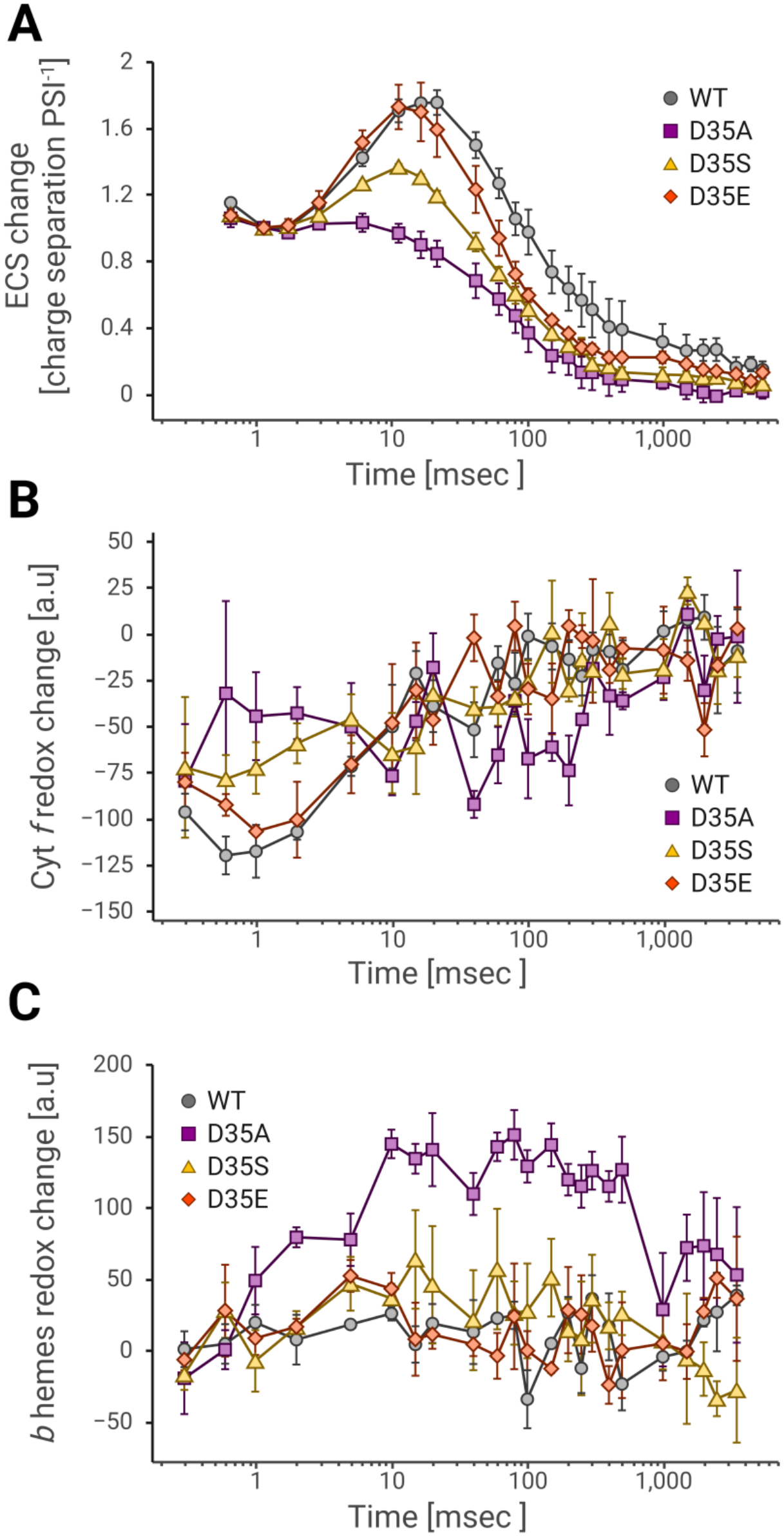
Flash-induced single-turnover redox absorbance changes of *b*-hemes and heme *f*: Single turn-over redox changes, derived from the sub-saturating flash-induced ECS changes at either; (**a**) 520 nm, representing membranal potential formation, (**b**) 554 nm, representing heme *f* redox state and (**c**) 563 nm, representing the redox state of heme *b*. The results are shown for the WT (grey circles) and the respective PetD:D35 mutants: D35A (purple squares), D35S (yellow triangles) and D35E (orange diamonds). ECS traces were gathered for 5 s from anoxic cultures by deactivating PSII with inhibitors. Heme redox state traces were gathered over the course of 3500 ms, from untreated cells. Averaged redox curves were fitted to determine re-reduction half times (ExpDec1; t_½_; in ms). Figure assembly was generated using Biorender.com.

In order to directly probe internal *b*_*6*_*f* electron transfer, flash-induced single-turnover absorbance changes were measured for Cyt*f* (at 554 nm, Figure.5B) and the *b*-hemes (at 563 nm, Figure.5C). Following a sub-saturating laser flash, wild-type complexes showed rapid oxidation of heme *f* within the first millisecond, followed by a re-reduction phase with a half-time of approximately 6 ms in accordance with past measurements (Buchert et al., 2020). Here again, D35E displayed WT-like kinetics (6.8 ms apparent half-time variation within error limits), while D35S showed a mid-range kinetics. Conversely, D35A showed severely disturbed heme *f* re-reduction kinetics, indicative of a redox-inactive or drastically hindered high-potential chain. Concurrently, single-turnover *b*-heme redox kinetics revealed that the low-potential *b*-heme remain significantly more reduced over an extended time window (10-500 msec) in the D35A variant. This stabilization of reduced *b*-hemes indicates a kinetic blockade at the Q_n_ site, impeding the forward electron transfer from heme *b*_*p*_ through heme *b*_*n*_ and *c*_*n*_, which secondarily restricts the high-potential chain and overall Q-cycle efficiency.

### EPR spectra of the hemes and the Rieske cluster

To gain more insights on the state of the hemes in the Q_n_ site, we recorded the EPR spectra over a broader magnetic field range, covering signals from the spin-spin-coupled hemes *b*_*n*_ and *c*_*n*_, through the high-spin and low-spin hemes, to the [2Fe-2S] cluster (Baymann et al., 2007; Szwalec et al., 2022). For all variants of air-oxidized *b*_*6*_*f*, the EPR transitions are nearly identical to those observed in the wild-type complex, suggesting that the introduced mutations do not significantly affect the proper assembly of the complex or the incorporation of its heme cofactors (Figure. 6). In the ascorbate-reduced samples, the g=3.49 transition of heme *f* disappears, while signals originating from the iron-sulfur cluster become apparent in the region below g=2.1 (Figure. 6A, supplementary Fig.S5). In all variants, the g=12 transition is clearly detectable, indicating the presence of spin-spin coupling between hemes *c*_*n*_ and *b*_*n*_ (Figure. 6B). The presence of transition at g=6, typically associated with an axial high-spin heme, suggests that there is a portion of axial high-spin heme which can be attributed to heme *c*_*n*_that is not spin-coupled to heme *b*_*n*_. A closer examination of this region in all the samples reveals a small increase in the linewidth of this transition in D35A and, to a lesser extent, in D35S, comparing to WT suggesting a slight deviation from axial symmetry towards a more rhombic geometry of the high-spin heme in these two mutants (Figure. 6B). Additionally, comparison of the signal amplitudes at g=12, g=6, and in the [2Fe2S] region suggests that the cofactor stoichiometry remains largely unchanged across all variants (supplementary Fig.S5). Overall, the EPR spectra of WT and the mutants confirm proper assembly and integrity of all cofactors *b*_*6*_*f* in mutated complexes.

**Figure 6.**
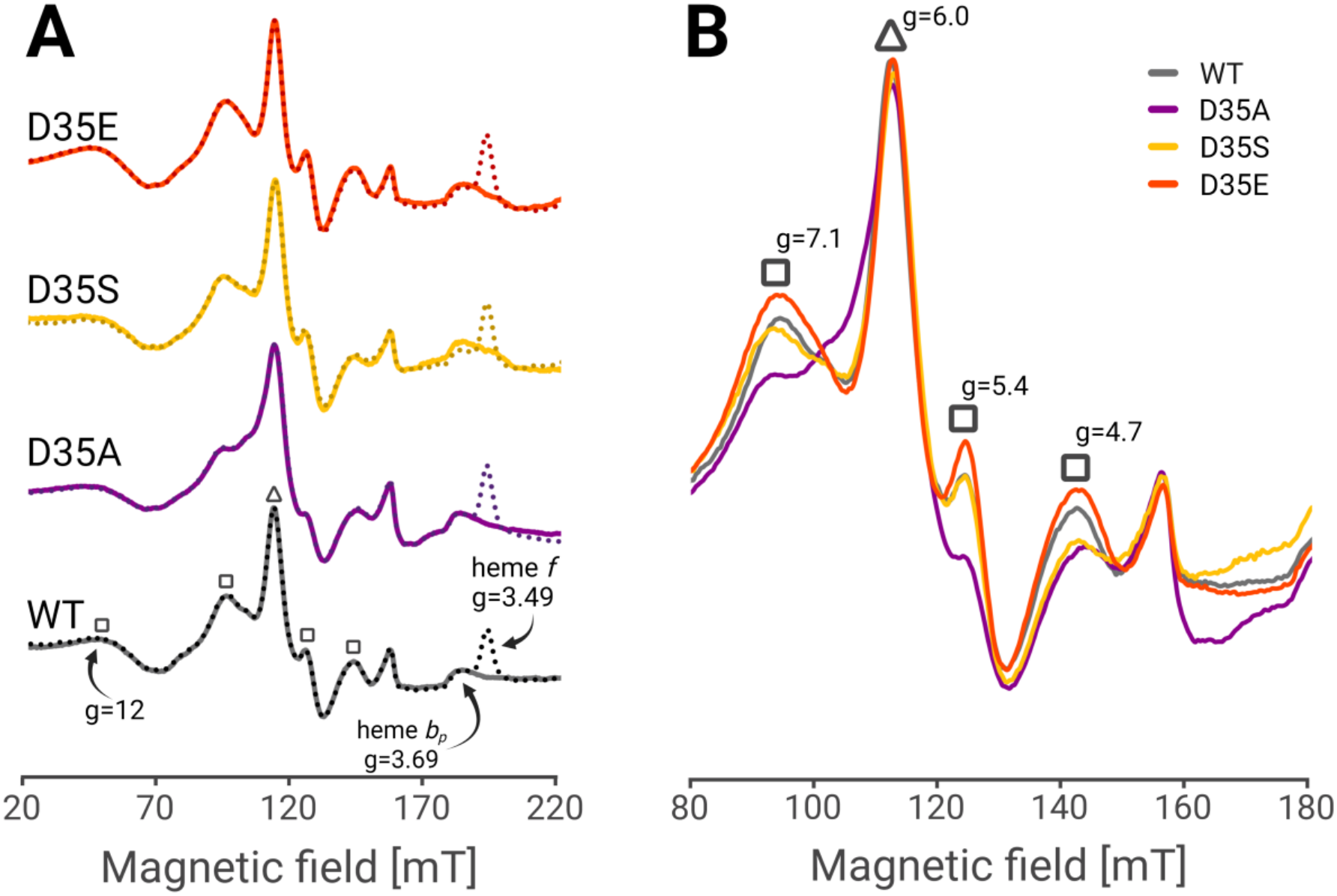
Low-temperature (10 K) X-band CW-EPR high-spin regions of isolated cytochrome *b*_*6*_*f* complexes: (**A**) Spectra compare untreated samples (solid lines) with ascorbate-reduced samples (dotted lines), for WT (gray) and PetD:D35 mutants: D35A (purple), D35S (yellow) and D35E (orange). Highlighted are the g-value benchmarks for heme *f* (g=3.49) and heme *b*_*p*_ (g=3.69) across WT. (**B**) Overlayed comparison of the strains between 80-180 mT, focusing on the transitions originating from heme *c*_*n*_-*b*_*n*_ coupled (□) and uncoupled (Δ) state, highlighting the peaks at g=12 and g=6.

## Discussion

The structural architecture of the stromal Q_n_ site in the *b*_*6*_*f* complex features unique specializations, most notably the heme *c*_*n*_/*b*_*n*_ pair, and an intricate network of conserved water molecules. In this study, we investigated the functional significance of PetD:D35, which anchors a key structural water molecule (W_2_) and forms the focal point of a putative proton resupply channel directed toward the plastoquinone (PQ) binding pocket. By generating and characterizing site-directed mutants (D35E, D35S and D35A) in *C. reinhardtii*, we provide direct *in vivo* and biophysical evidence that this residue is essential for efficient Q-cycle turnover, proton translocation, and overall photosynthetic viability. Biochemical evaluations and low-temperature EPR spectroscopy confirmed that substituting PetD:D35 does not destabilize the complex assembly or disrupt the intrinsic coordination of the Rieske [2Fe-2S] cluster and high-spin hemes in isolated dimeric *b*_*6*_*f* complexes (Figure.2). However, flash-induced single-turnover kinetics revealed a severe functional bottleneck. While the conservative D35E substitution, in which the carboxylic acid group is retained, exhibited wild-type-like kinetics and growth phenotypes, the removal of the negative charge and proton-donating capability crippled the complex, as was the case in the D35S mutant. Moreover, when this residue loses all its ability to bridge out the water channel (as in D35A), the complex begins to lose its activity, and the culture cease to grow in autotrophic conditions. Specifically, single-turnover absorbance changes at 563 nm demonstrated that low-potential *b*-hemes remain significantly reduced over extended time windows (10-500 msec, Figure.5C). This stabilization of reduced *b*-hemes points directly to a kinetic blockade at the Q_n_ site. In the absence of the carboxylate group and its associated water network (Figure.1 C,D), proton delivery to the bound PQ is impaired, limiting its two-electron reduction to PQH_2_ at the Q_i_ site. This restricts forward electron transfer through the low-potential chain, thereby limiting PQH_2_ oxidation at the Q_0_ site and, consequently, the re-reduction of heme f (Fig. 5B).

The impaired catalysis at *b*_*6*_*f* exerts profound effects on the reduction state of the PQ pool. Chlorophyll *a* fluorescence measurement revealed early-stage drop in the apparent quantum yield of PSII for D35S and to a greater extent, D35A (Figure.3B). Moreover, ETR assessments showed an over-all lower flow rate for D35A under steady state illumination, both under aerobic and anoxic conditions (Figure.3 C,D). Downstream to *b*_*6*_*f*, P700 kinetics showed an elevated fraction of constantly oxidized P700^+^ (Y_ND_) under steady-state illumination, reflecting a severe donor-side limitation caused by sluggish plastocyanin resupply from the hindered complex (Figure.4). On top of that, electrochromic shift (ECS) measurements further illuminated these defects: while WT and D35E cells showed robust b-phase development reflecting normal proton motive force generation and Q-cycle activity, the D35S variant lacked a distinct b-phase increase, and the D35A was completely absent of it (Figure.5A). This confirms that the proton-resupply pathway mediated by the water channel that runs through PetD is critical not only for chemical reduction of PQ but also for maintaining a functional transmembrane proton gradient. Interestingly, this overall diminished activity had a negligible (if any) effect on light harvesting state-transitions (supplementary Fig.S4). This process is controlled by kinases such as state transition-deficient mutant 7 (STT7) (Depège et al., 2003), which were shown to be responsible for this distribution of energy between the photosystems (Bergner et al., 2015; Nawrocki et al., 2016). Indeed there is a link between state transition and *b*_*6*_*f* availability and *b*_*6*_*f* dependent electron transfer (Zito et al., 1999; Zaeem et al., 2026). In addition, there is evidence that STT7 and *b*_*6*_*f* physically interact with each other (Shapiguzov et al., 2016; Dumas et al., 2017; Riché et al., 2024; Zaeem et al., 2026). Despite, the strong impact in *b*_*6*_*f* dependent electron transfer in D35A (Figure.4, 5), state transitions occur in this strain (Supplemental Fig.S4). Therefore, interaction between STT7 and *b*_*6*_*f* appears to be intact and activation of the STT7 kinase in place. It has been suggested that the function of the Q_p_-site is required for the activation of STT7 (Vener et al., 1997; Zito et al., 1999). Our data indicate, that impaired Q_n_ function still allows state transitions, indicative for STT7 kinase activation, despite a severe slow-down of electron transfer at Q_p_. It is thus tempting to speculate, that mutation of the conserved PEWY motif in the EF loop of PetD into PWYE (Zito et al., 1999) interferes with STT7 binding and activation, rather than STT7 activation being directly dependent to the electron transfer activity at Q_p_ site. It has been shown that the activity of STT7 is linked to a feed-back inhibition of the kinase via phosphorylation of PetD T4 (Zaeem et al., 2026). Whether the electron activity of *b*_*6*_*f* is linked to STT7 activation remains to be re-investigated.

## Materials and Methods

### Plasmid design and Transformation

Plasmids containing the mutated petD gene as well as the aadA cassette (Goldschmidt Clermont, 1991; hereafter termed ‘cassette’) as a selectable marker for spectinomycin resistance were created with common cloning approaches and directed primer design. Using the pUC18-3’-petD-aadA-5’ vector (Zaeem et al., 2026) as a template, PCR was performed using primers 1 and phosphorylated primers 2 – 4 respectively. The purified products were ligated to the fragment, gained from PCR with primers 5 and 6 to introduce three individual point muta ons into the Chlamydomonas petD gene (petD D35E, D35S and D35A). The resulting fragment and vector were digested (PacI/SalI) and swapped into the pUC18-3’-petD aadA-5’ vector. For PCR amplification either the Phusion™ DNA polymerase in combination with the High Fidelity (HF) buffer (both Thermo Fisher Scientific) or the Phusion™ Plus Green PCR-Master mix (Thermo Fisher Scientific) were used. PCR was performed with self-optimized Thermocycler programs (not shown), using the Mastercycler nexus GX2 (Merck KGaA). For ligation the T4-DNA ligase in combination with the 10x T4-DNA ligase buffer (both Thermo fisher scientific) were used. Liga on was performed for 1 hour at 37 C. Plasmids were purified from E. coli cells, using the NucleoSpin Gel and PCR Clean-up XS kit from Macherey-Nagel. For DNA fragment purification, the NucleoSpin Plasmid EasyPure kit from Macherey-Nagel was used. Escherischia coli NEB5α cells as recipients were thawed on ice. LB plates with ampicillin (100 μg/ml) were prepared. Just before the heat shock, the mix containing the ligated plasmids was added to the cells and le on ice for 10 minutes. A er heat-shocking the cells at 42 C for 32 seconds, cells were recovered with warm LB and plated out to grow at 37 C over night.

### Strains and conditions

The *Chlamydomonas reinhardtii* strains used in this work are derived from the ma ng-type plus (mt+) 137C background containing a C-terminal 6xHis-tag at the *b*_*6*_*f* subunit PetA (Choquet et al., 2003). Cells were grown at ∼50 µE m^-2^ s^-1^ with 16-h light/8-h dark cycles at 23 C in either Tris-acetate-phosphate (TAP) medium or in Tris-phosphate (TP) minimal medium (Gorman & Levine, 1965; Tab. 3). Optionally, 1.5 % agarose was added. For selective TAP and 28 Methods TP media 150 µg/ml Spectinomycin was used (TAPspec). Cultures for physiological measurements (PAM and JTS) were grown in TP and aerated by constant bubbling with sterile air. Cultures for biochemical procedures, biolistic transformation, 77K chlorophyll fluorescence measurements and phototrophic growth tests were grown in TAP and all cultures were diluted the day before the experiment and measured in their exponential growth phase.

### Directed mutagenesis

#### Biolistic Transformation

Tungsten microcarriers M-10 (0.7 µm) were boiled in absolute ethanol at 90 C for 2 hours and washed with sterile ddH2O before storing them in 50 % glycerol (w/v) at - 20 C for no longer than two weeks. Per plate 1 mg Tungsten particles were taken. To coat the par cles with the desired plasmid, 0.5 μg of DNA, 1 M CaCl2 and 16 mM spermidine were added. The mix was kept on ice for 0.5 hours and vortexed vigorously every few minutes. Biolistic transformation of the wild-type recipient strain was performed, using the PDS 1000/He Biolistic Particle delivery system (Bio-Rad). Beforehand, 4 x 107 cells/plate were streaked out on TAPspec plates and biolistic transformation was carried out according to the manufacturer’s manual. Afterwards, the plates were incubated at ∼50 μmol photons m-2 s-1 for about 2 weeks un l colonies appeared

#### Mutant screening

After about 2 weeks of growth on selective TAP plates the grown colonies were picked, and re-streaked on TAPspec several times. To screen the mutant colonies for homoplasy, DNA was extracted using the dilution buffer from Thermo Fisher Scientific and used as a template for the Screening-PCR with primers 5 and 6 (Tab. 5). Homoplasmic clones show the cassette containing fragment at 3477 bp, while the wild-type gene is at around ∼2000 bp. The homoplasmic region of positive clones was amplified by PCR (with primers 6 and 7) and sent for Sequencing with primers 6 and 9, using the Eurofins genomics ‘Tube seq supreme’ service.

### Biochemical methods

#### Thylakoid isolation

Thylakoid isolation was performed according to the PSI isolation procedure described in (Naschberger et al., 2022; Ogawa et al., 2026) with some minor modifications. Cells were grown under aerobic conditions and cell harvest, thylakoid extraction and separation by 3-step sucrose density gradient (5 mM HEPES/KOH, pH 7.5, 10 mM EDTA and sucrose concentrations of 0.33, 1.3 and 1.8 M), Thylakoid membranes were then extracted using a Pasteur pipette, set to a chlorophyll concentration of 0.5 mg Chl ml^-1^ and stored in the lowest density 3-step gradient buffer at -20 C.

#### Cytochrome b_6_f isolation

Methods For the *b*_*6*_*f* isolation, the PetA 6xHis tagged strains were grown in 5 L of TAP under aerobic conditions and harvested in the exponential growth phase. The complex preparation was done according to (Stroebel et al., 2003) with modifications as described in (Milrad et al., 2025). Membrane solubilization was performed using 0.5-0.8 % (w/v) 4-trans-(4-trans-propyl cyclohexyl)-cyclohexyl-α-maltoside (tPCCαM) for 45 minutes on ice. Insolubilized thylakoids were pelleted down by centrifuging the sample at 50,000 g for 15 min. Supernatant was loaded on NiNTA columns (5 mM HEPES/KOH, pH 7.8, 100 mM KCl 5 mM Imidazole and 0.01% tPCCαM). Elutants were filtered using Amicon® ultra size exclusion filter tube (Merck KGaA, 100.000 MW) and loaded on a linear sucrose gradient (5 mM HEPES/KOH, pH 7.8, 5 mM

MgCl_2_, 5 mM CaCl_2_,0.7 mM Sucrose and 0.01% tPCCαM, 10 mL total in SW41-tubes, 200,000 g for ∼16 hr) chromatography and sucrose density gradient centrifugation were performed. The brown frac on was extracted using a Pasteur-pipette and an Amicon® ultra size exclusion filter tube (Merck KGaA, 100.000 MW) was used to change the buffer for storage at -80 C (30 mM MOPS, pH 7.0, 0.01 % tPCCαM, 30 mM KCl) and to determine protein concentration. Spectra of reduced proteins were measured from 350-750 nm by adding ∼10 mM ascorbate. The absorption of reduced cytochrome *f* at 554 nm was baseline corrected (baseline drawn from 543 and 560 nm) and used to calculate the protein concentration according to Lambert-Beer (ε554 = 24 mM cm^-1^ for *C. reinhardtii* cytochrome *f*).

#### SDS PAGE and Blotting

Isolated proteins and thylakoid membranes were boiled in 1x sample loading buffer after adding 166 mM Na_2_CO_3_ and 166 mM dithiothreitol (DTT) for 15 minutes. For PetD expression, 5 µgr Chl from isolated thylakoids were taken (see section 5.4.1). The heme staining gel was loaded with 10 µgr (= 100 %) isolated complexes, that were incubated without DTT and at 37 C. For the separation of proteins, a 13 % (w/v) SDS PAGE was performed (Schägger and Von Jagow, 1987). Transfer onto a nitrocellulose membrane (Amersham™ Protran®, Merck KGaA) was done with Whatman paper (3MM, Merck KGaA) soaked in different buffers on the anode-(25 mM Tris-HCl, pH 10.4, 20 % isopropanol (v/v)) and cathode-facing side (25 mM Tris-HCl, pH 9.4, 20 % isopropanol (v/v), 40 mM ACA, 0.001 % SDS (w/v)) of the gel-membrane stack. For Ponceau-staining the membrane was incubated with 0.2 % Ponceau S and 3 % TCA in ddH_2_O before blocking (3,75 % milk powder (w/v)). Blocking and all the following washing steps were performed using 1x PBS buffer (137 mM NaCl, 2.5 mM KCl, 40 mM Na_2_HPO_4_, 15 mM KH_2_PO_4_) with 1 % Tween. Primary antibody incubation was done for 1 hr with a PetD antibody (gift from FA Wollmann, IBCP Paris) and AtpB antibody (Agrisera). For visualization a secondary An-IgG (Rabbit)-Peroxidase antibody (Invitrogen) was used in combination with the ECL solutions from Merck KGaA (Immobilon ECL Ultra-Western-HRP-Substrate) on the Fusion SL (Viber Lourmat) imager. To visualize the presence of high-spin hemes, 10 µg isolated *b*_*6*_*f* complex were stained in a non-reducing SDS-PAGE gel by addition of 3,3’,5,5’-tetramethylbenzidine (TMBZ) according to (Thomas and Poznansky, 1990). Pictures were taken a er adding 30 mM H_2_O_2_.

### Physiological measurements Methods

#### Joliot-type-spectroscopy

For optical spectroscopy a Joliot-type spectrophotometer (JTS-150, SpectroLogiX) was used to measure the electrochromic shift (ECS) that depends on changes in the transmembrane electric field induced for example when pmf is generated. The ECS signal is expressed according to absorbance intensity for reference and sample probe as ΔI/I and detecting photodiodes were protected from scattering ac nic light by applying 3 mm BG39 (Scho, Mainz, Germany) filters for ECS from carotenoid absorption and P700 filters (Scho, Mainz, Germany) for ECS derived mainly from chlorophylls. Electron transfer rates (ETR), P700 kinetics (P700) and *b*_*6*_*f* heme redox kinetics were measured according to (Buchert et al., 2020; Zaeem et al., 2026). Chlorophyll concentration was set to 20 µgr Chl ml^-1^ in 20% TP Ficoll (w/v), the cells were kept in the dark for ∼10 minutes. By applying 2.5 sec actinic light / 2.5 sec dark intervals (630 nm LED, 600 µE) over the course of ∼1 minute, the cells were adapted to the short 250 µsec dark intervals required for the ECS detection light pulses (10 µsec after 200 µsec). To measure ETR, the a-phase (see Bailleul et al., 2010) was normalized to PSI by measuring initial ECS rise induced by a saturating 6 ns laser flash (Q-switched Nd:YAG, Continuum) before and after the addition of 20 µM DCMU and 2 mM HA to inhibit PSII contribution. Dark-interval relaxation kinetics (DIRK) measurements were used to determine the steady-state ETR in charge separations PSI^-1^ s^-1^ (Sacksteder and Kramer, 2000; Joliot and Joliot, 2002; Zaeem et al., 2026). Using the same PSII inhibition approach, the P700 kinetics were measured according to (Alric et al., 2009) and used to quantify the amount of oxidized P700^+^ over the course of 2.5 s actinic illumination (630 nm LED, 600 µE m^-2^ s^-1^) and 2.5 sec darkness. The 20 msec saturating light pulse (630 nm LED, 3000 µE m^-2^ s^-1^) was applied before transitioning to darkness and the determined value in the presence of HA and DCMU was used for normalization to the relative concentration of oxidized PSI (P700^+^). The same measurements were performed under anoxic conditions. To induce anoxic conditions a mix containing 30 U ml^-1^ Catalase (Sigma Aldrich, Germany), 10 U ml^-1^ glucose oxidase (Carl Roth GmbH) and 50 mM glucose was added and incubated in the dark for 30 minutes. For the single turnover, flash-induced *b*_*6*_*f* heme redox measurements, the cell concentration of aerobic cultures in the exponential growth phase was set to 40 µg Chl ml^-1^ and the JTS-150 in combination with a legacy white light LED lamp and interference filters (520 nm, 546 nm, 554 nm, 563 nm and 573 nm) was used. Cultures were adapted to darkness for 10 minutes and single turn-over sub-saturating (∼50 %) 6-ns flashes (Q-switched Nd:YAG, Continuum) were applied a er 3 s of darkness, followed by an additional 3.5 s of darkness. For baseline correction, three intervals without applying a flash were averaged and subtracted. Five technical replicates were performed for each wavelength. Wavelength 554 nm and 563 nm were measured for heme *f* and the *b* hemes respectively with correction wavelength measurements at 546 nm and 573 nm. Deconvolution procedure and weigh ng of the correction wavelength for the different types of hemes were described in (Buchert et al., 2020; Zaeem et al., 2026). Methods Based on the normalized saturating light pulse (Q-switched Nd:YAG, Continuum) ECS changes under anoxic conditions with PSII inhibitors, a-phase (PSI contribution), b-phase (*b*_*6*_*f* contribution) and c-phase (*b*_*6*_*f* contribution and ATP-synthase consumption) can be visualized (Bailleul et al., 2010). By fi ng the respective c-phase with the ExpDec2 function (shown below) and subtracting it from the averaged ECS signal, the corrected b-phases (*b*_*6*_*f* contribution to ECS build-up) were obtained.

#### Photoautotrophic growth tests

To assess photoautotrophic growth, TAP grown cells were set to either 106 cells/mL or to 2.5 µgr Chl ml^-1^ with TP medium. For each biological replicate 4 technical replicates of 40 μL (and 20 µL) cell suspension were dropped onto TP agar plates and incubated at low-light (∼50 µE m^-2^ s^-1^) and high-light (∼200 µE m^-2^ s^-1^) conditions for 6-9 days. For anoxic conditions, the GENbox anaer anoxia bags (bioMerieux) were used.

#### Pulsed-amplitude modulation (PAM) fluorometry

Pulsed-amplitude chlorophyll fluorescence modulation (PAM) fluorometry was performed using the Maxi-Imaging PAM chlorophyll fluorometer (IMAG-MAX/K2, Walz Germany). The chlorophyll concentration of TP cells in the exponential growth phase was set to 5 μgr Chl ml^-1^ and cultures were adapted in the dark for 20 minutes before the experiment. First measuring F0 in the dark and subtracting it from Fm, measured a er a 250 msec saturating pulse (520 nm LED, 3000), Fv=Fm–Fo can be estimated. By repeatedly measuring Fm (Fm’), the PSII quantum yield (Fv’/Fm’ or (Fm’-Fo’)/Fm’) and NPQ ((Fm – Fm’)/Fm’) can be estimated over me and with background illumination ((Maxwell and Johnson, 2000; Oxborough, 2004). Calculations were generally performed with the Walz software (see Walz manual; IMAGING-PAM MAXI-version). Additionally, the reduction state of PSII (or the fraction of open PSII reaction centres) was determined based on the puddle model of PSII antenna pigment organization with qP defined as (Fm’ – Fo)/(Fm’ – Fo’) and plotted as 1-qP at increasing background light intensities. In this work, two background ac nic light (AL) illumination protocols were used that (i) apply constant AL illumination of ∼280 µE for 10 minutes, followed by 10 minutes of darkness and (ii) an increasing intensity of AL illumination of ∼10 to ∼700 µE over the course of around 5 min.

#### 77 K chlorophyll fluorescence

For the assessment of state transitions chlorophyll fluorescence spectra were measured with the JASCO spectrofluorometer (FP-6500 series). At a chlorophyll concentration of 5 µg Chl ml^-1^ different treatments were applied to evoke the desired state I and state II conditions. For state I induction, the cells were le shaking at 50 µE m^-2^ s^-1^ for 20 min with 10 µM DCMU. To induce state II conditions 10 mM glucose and 50 U mL^-1^ glucose oxidase (Sigma Aldrich, Germany) were added, and the cells were incubated in the dark for 20 min. Right a er the treatment cell aliquots were frozen in liquid nitrogen and spectra from 600 – 800 nm were measured with three technical replicates per sample. The excitation wavelength was 436 nm (5 nm band width) and samples were measured at 77K in liquid nitrogen. Data was normalized to the peak at 687 nm, that is attributed to the PSII contribution.

### EPR measurements

EPR spectra were recorded using a Bruker ELEXSYS E580 spectrometer equipped with an Oxford Instruments ESP900 cryostat inserted into an SHQ4122 resonator. Low-temperature conditions were achieved using a ColdEdge closed-cycle helium cryocooler (Stinger) system, with temperature regulation provided by a Lake Shore 350 temperature controller. Heme spectra were recorded at 10 K, whereas spectra of the [2Fe–2S] cluster were acquired at 20 K. The measurement parameters for heme signals were as follows: modulation amplitude, 15 G; microwave power, 6 mW; conversion time, 40.96 ms; time constant, 10.24 ms; and sweep time, 41.94 s. For the [2Fe–2S] cluster, the following parameters were used: modulation amplitude, 10 G; microwave power, 2 mW; conversion time, 40.96 ms; time constant, 10 ms; and sweep time, 41.94 s.

## Supplementary Figures

**Supplementary Figure S1:**
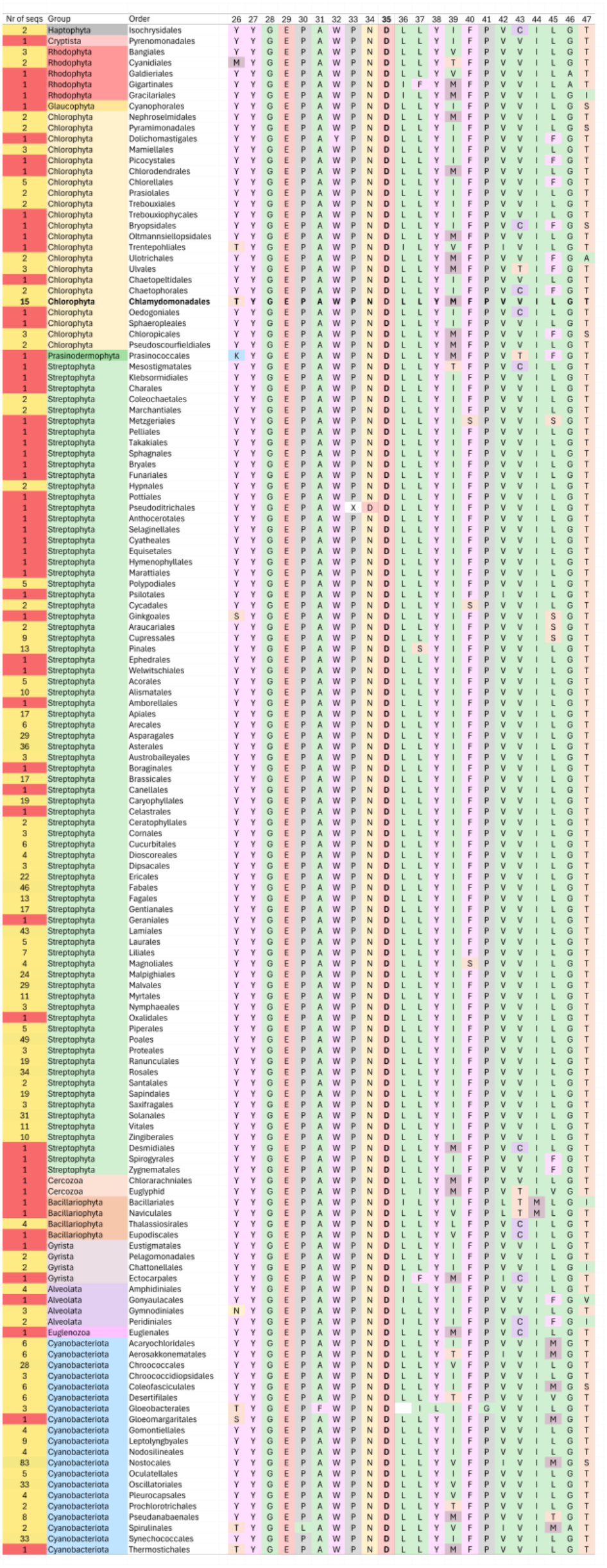
Loop sequence alignment of for PetD. Sequence alignments were conducted by taking all available sequences from Uniprot.org, using the keyword: PETD. In total, 976 sequences had a structural similarity from which all organisms were clustered based on their order and aligned according to conserved regions, with the highest similarity. Number of sequences in each group is indicated, and orders are clustered to phylums. Over-all, aspartate 35 is featured to be highly conserved throughout the entire photosynthetic kingdom.

**Supplementary Figure S2:**
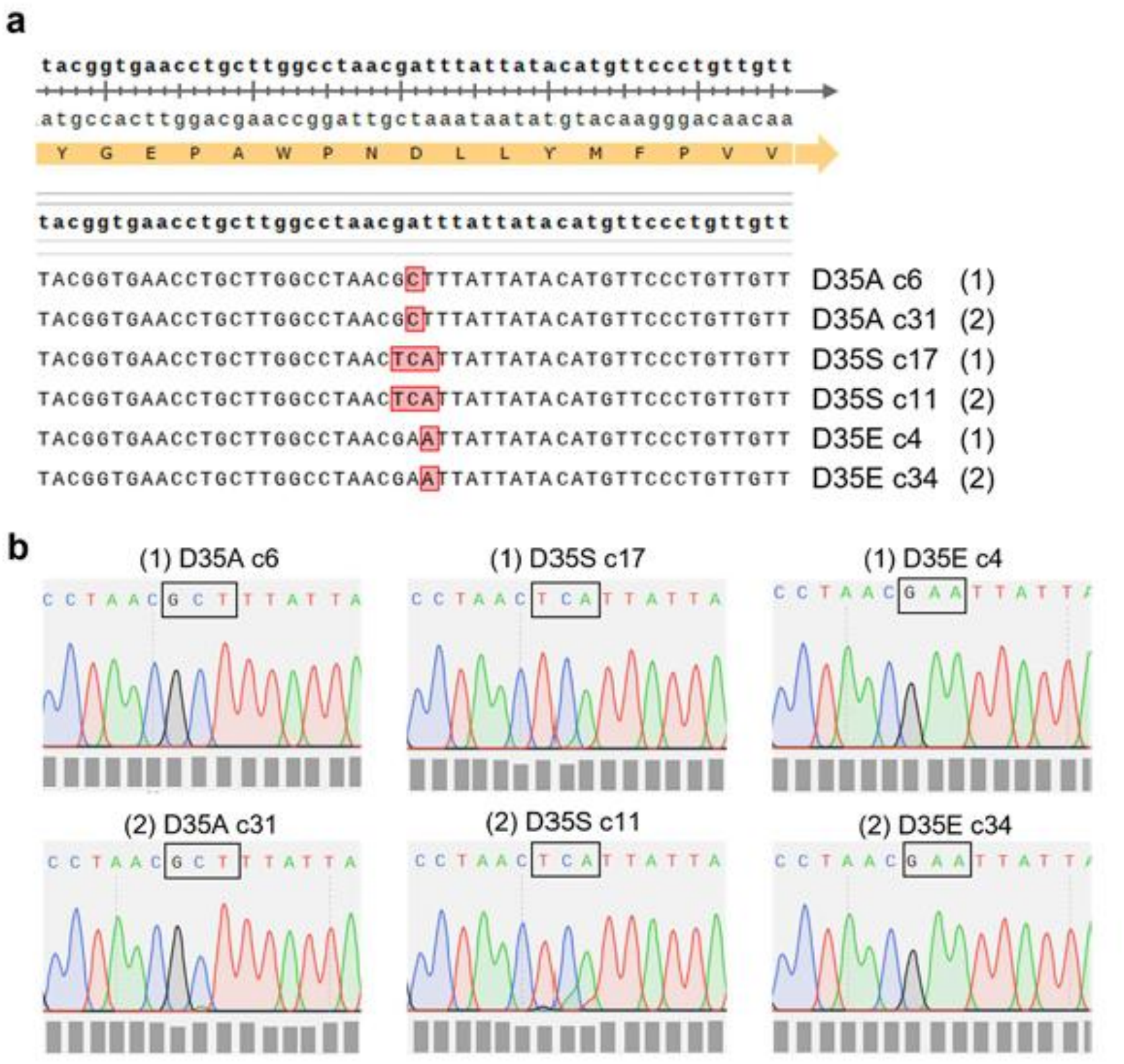
Sequencing results for the generated substitution mutants. Purified PCR products were sent, premixed, with the primer: ctcctaattaattaattcccattttatactg. (**a**) Results for two homoplasmic clones of theSubIV:D35 mutants D35A (c6 and c31), D35S (c17 and c11) and D35E (c4 and c34) were aligned to the native PetD gene as part of the pUC18-3’-petD-aadA-5’ vector using Snapgene. (**b**) Corresponding chromatograms with the quality values determined by Snapgene (grey boxes). The desired codon substitution is indicated (black box). Clones (1) were chosen for mutant characterization unless stated otherwise.

**Supplementary Figure S3:**
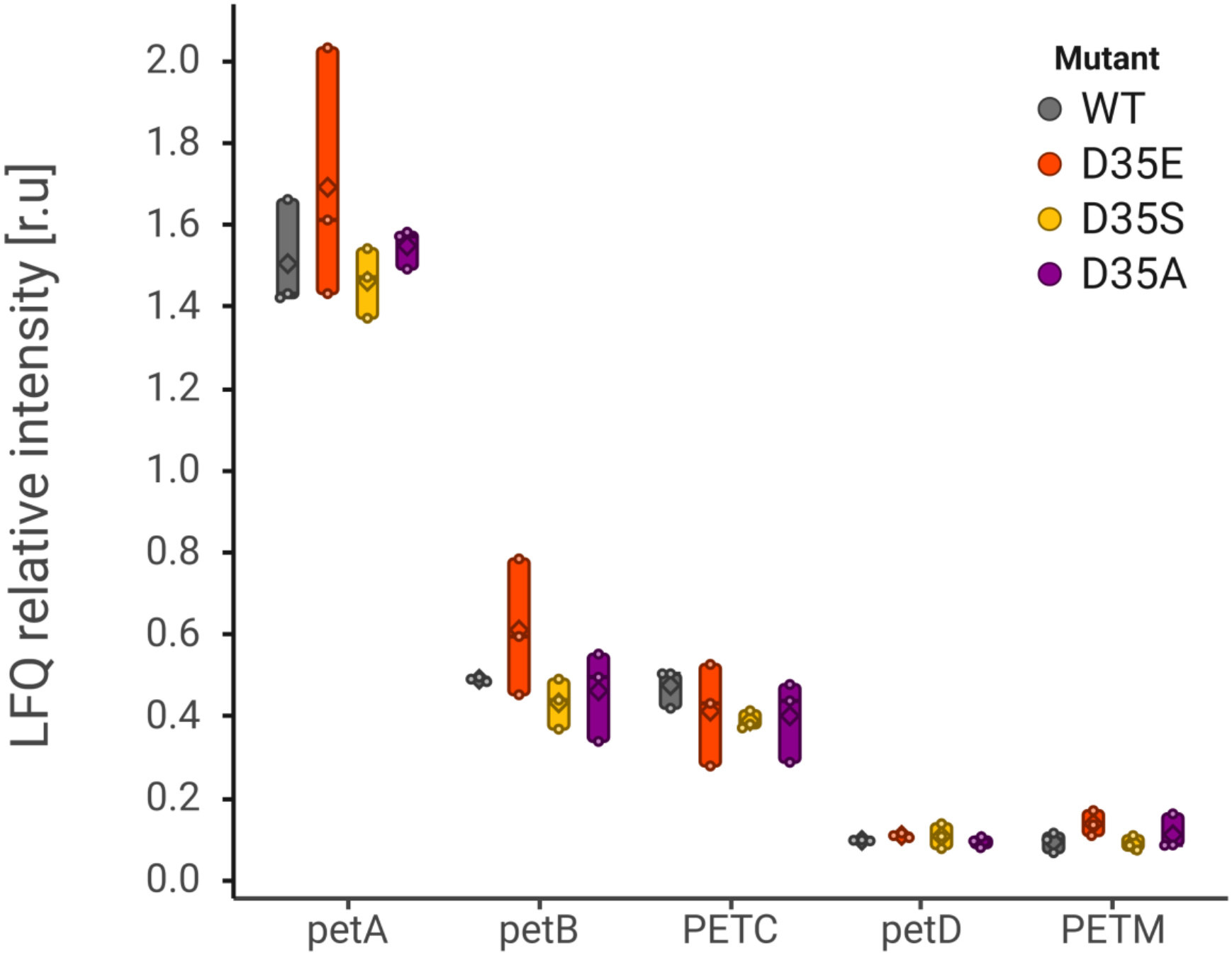
Gene expression for Cyt*b*_*6*_*f* subunits. All strains were grown in TAP medium at 25 C in normal light conditions with light intensities in 60 µE m^-2^ s^-1^. Relative LFQ intensities of petA, petB, PETC, petD and PETM protein abundances are expressed as boxplots from three independent biological replicates. Normalization was conducted in relation to the expression of ATPase subunits subunit (an average of; atpA, atpB, ATPC, ATPD, atpE and atpF). Figure was assembled using Biorender.com.

**Supplementary Figure S4:**
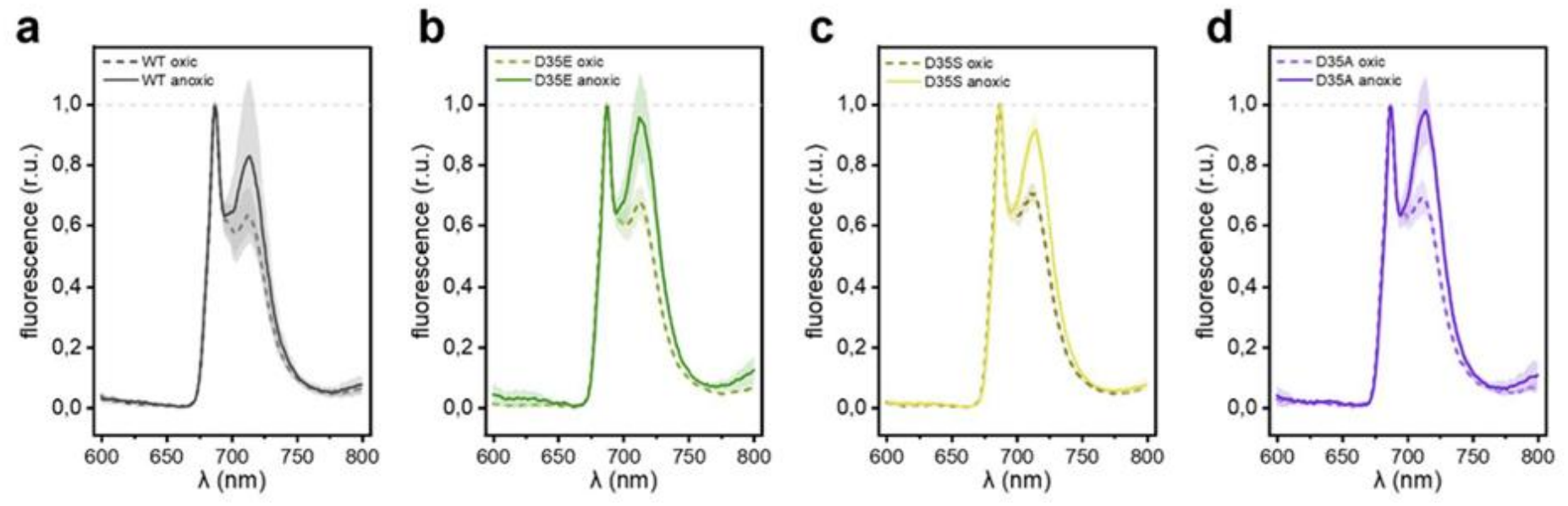
Averaged relative fluorescence spectra at 77 K. Three biological replicates were normalized to PSII fluorescence contribution (687 nm) for (a) WT and (b–c) the D35sIV mutant strains D35E, D35S and D35A after artificial induction of state II (anoxic) and state I (dashed line; oxic) by reducing and oxidizing the PQ pool, respectively. Cultures were grown heterotrophically.

**Supplementary Figure S5:**
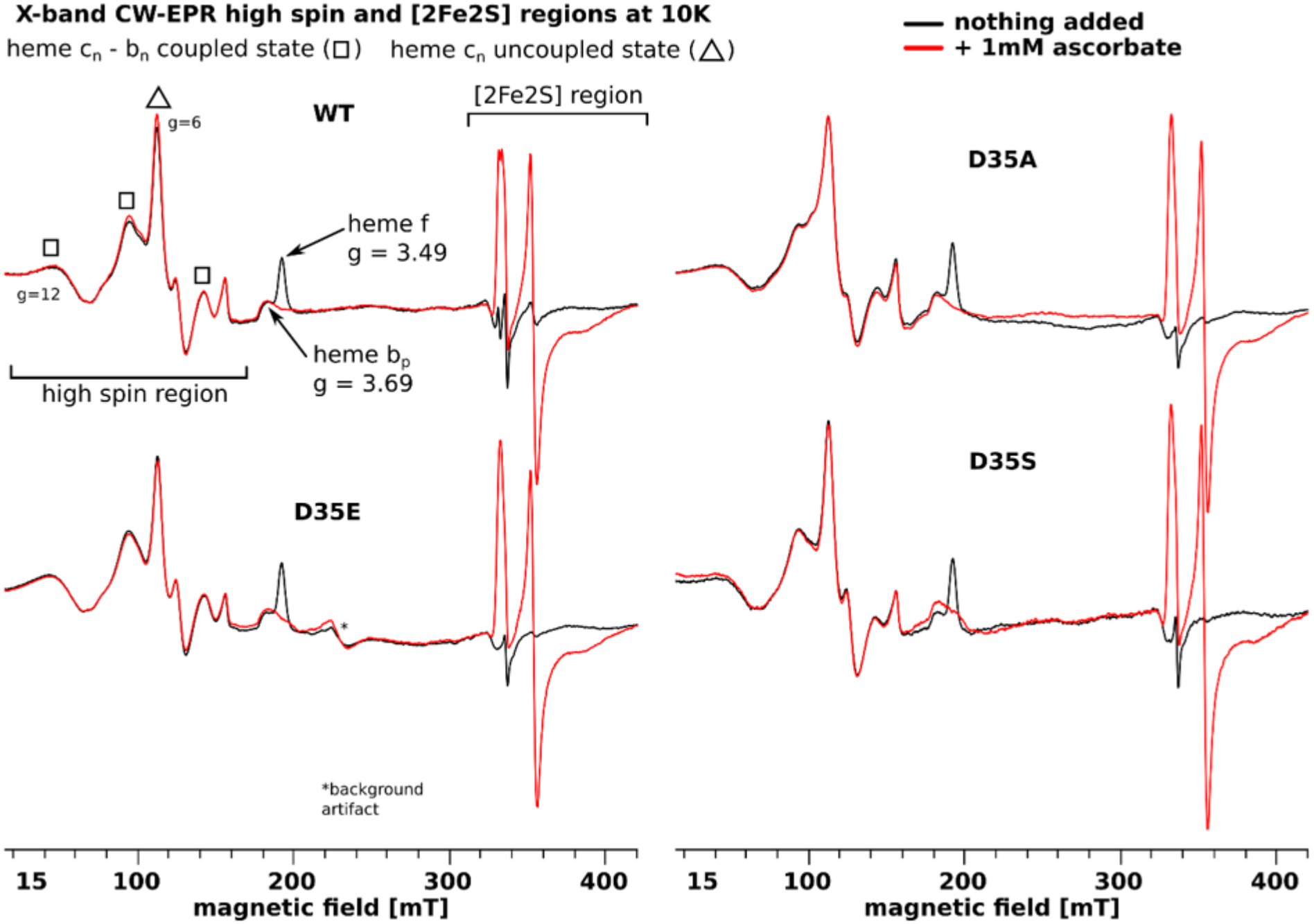
Full EPR spectra. Spectra compare untreated samples (black lines) with ascorbate-reduced samples (red lines), highlighting the *c*_*n*_-*b*_*n*_ coupled state (□) and uncoupled state (Δ) alongside g-value benchmarks for heme *f* (g=3.49) and heme *b*_*p*_ (g=3.69) across WT (**a**) and PetD:D35 mutants: (**b**) D35E, (**c**) D35A and (**d**) D35S.

